# A flexible, high-throughput system for studying mRNA translation kinetics *in vitro* and *in cellulo* with HiBit technology

**DOI:** 10.1101/2024.06.27.600987

**Authors:** C. Ascanelli, E. Lawrence, C. A. P. Batho, C. H. Wilson

**Affiliations:** Department of Pharmacology, University of Cambridge, 80 Tennis Court Road, Cambridge CB2 1PD, UK

## Abstract

HiBit is an engineered luciferase’s 11 amino acid component that can be introduced as a tag at either terminus of a protein of interest. When the LgBit component and a substrate are present, HiBit and LgBit dimerise forming a functional luciferase. The HiBit technology has been extensively used for high-throughput protein turnover studies in cells. Here, we have adapted the use of the HiBit technology to quantify mRNA translation temporally *in vitro* in the rabbit reticulocyte system and *in cellulo* in HEK293 cells constitutively expressing LgBit. The assay system can detect differences in Cap, 5’UTR, modified nucleotide composition, coding sequence optimisation and poly(A) length. Importantly, using these assays we established the optimal mRNA composition varied depending on the encoded protein of interest, highlighting the importance of screening methods tailored to the protein of interest, and not reliant on reporter proteins. Our findings demonstrated that HiBit can be easily and readily adapted to monitor mRNA translation and offers a novel and highly favourable method for the development of mRNA-based therapeutics.

**Graphical abstract:** 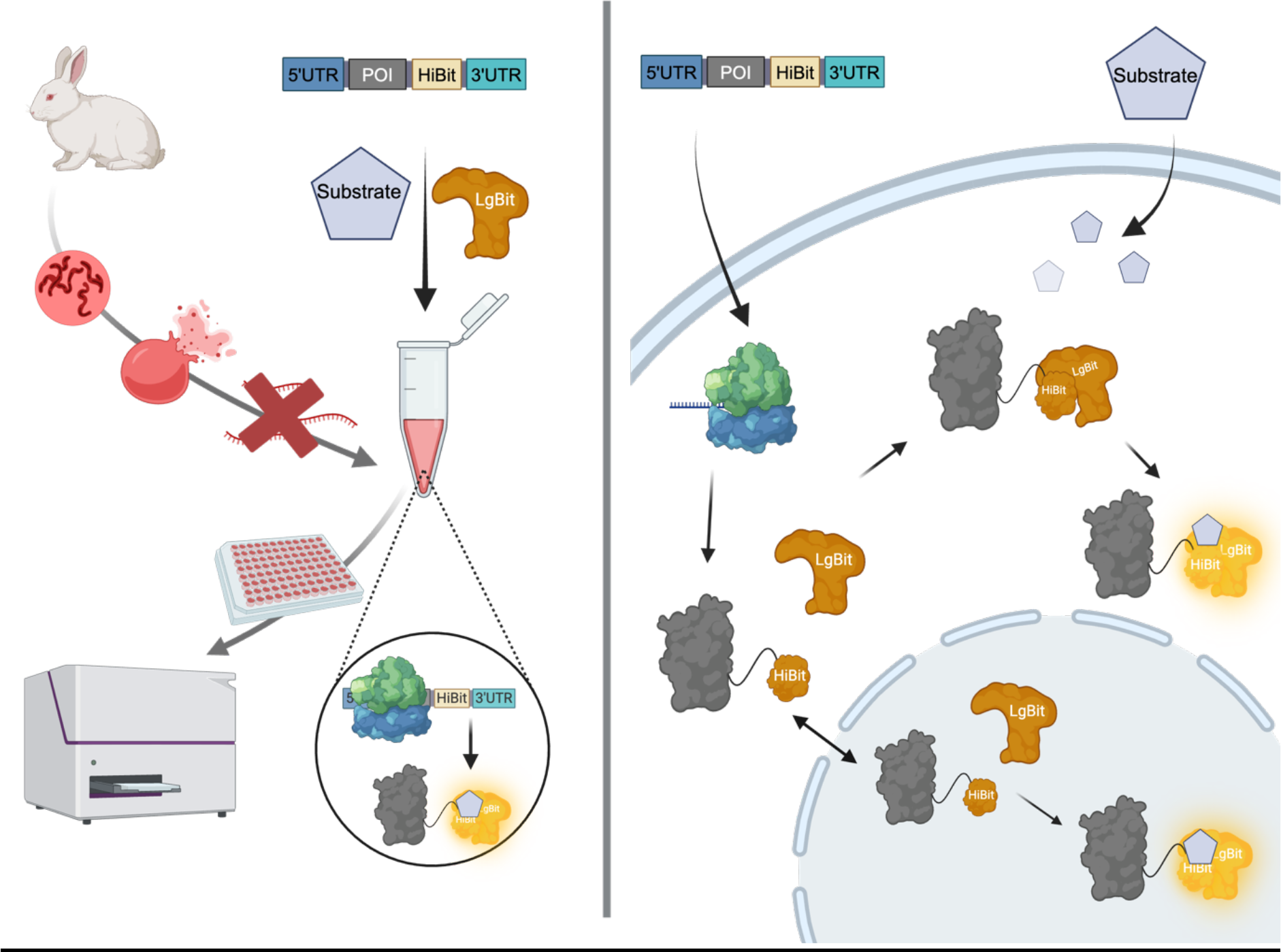

## Introduction

Fuelled by the successful development and worldwide roll-out of messenger RNA (mRNA) vaccines during the COVID-19 pandemic(1), novel therapies based on mRNA are a rapidly expanding field. In particular, therapeutic mRNAs are in development as vaccines for numerous infectious diseases and cancers, and as protein replacement therapeutics for genetic diseases, such as cystic fibrosis, autoimmune disorders, metabolic disorders and cardiovascular disease(1). The therapeutic modality is exceptionally versatile because mRNA can be engineered to express any peptide, protein, or group of proteins transiently, by harnessing the target cell’s endogenous protein synthesis apparatus. Messenger RNA therefore offers a wealth of advantages over DNA or protein/peptide drugs, such as higher transfection efficiency, lower toxicity(2), no infection risk, and a better safety profile than DNA-based therapies as it does not enter the nucleus and pose a risk of insertional mutagenesis(1, 3). Messenger RNA overcomes barriers that exogenously delivered amino acid-based therapeutics suffer such as incorrect/inefficient folding, assembly and post-translational modifications(4). Furthermore, the manufacture of mRNA is cell-free therefore its production is convenient, rapid, scalable, and low-cost. Given these remarkable characteristics, mRNA therapeutics offer a promising versatile prospect for drug developers.

The pipeline for the development of mRNA-based therapies consists of mRNA design and optimisation, synthesis and purification, entrapment in vehicles for delivery, pharmacokinetic and pharmacodynamic evaluation, manufacturing and clinical assessment(1). The design and optimisation stages are complex as many structural elements converge to yield an optimal mRNA-based therapeutic. A mature mRNA consists of the coding region, flanked at either end by untranslated regions (UTRs), a 7-methyl-guanylate (m^7^G) cap at the 5’ end which binds the eukaryotic translation initiation factors (eIFs), and a poly(A) tail which protects the mRNA from decay by deadenylation if bound by cytoplasmic poly(A) binding protein (PABPC)(5).

Messenger RNA optimisation is multifaceted, involving a range of techniques which can improve mRNA stability, translation efficiency, protein expression and overall enhance efficacy of mRNA-based therapies and vaccines. Design rules for the components of therapeutic mRNA are beginning to be established to optimise protein expression inside cells(6, 7). However, the design rules are often based upon highly engineered reporter proteins in which translation is extremely efficient, such as reporter proteins GFP(7) and NanoLuciferase(8) or SARS-CoV-2 spike protein(8, 9) and it is likely each therapeutic protein of interest (POI) in each cell of interest may have an optimum design which is not yet appreciated.

One pivotal aspect of mRNA optimisation is the design and engineering of the UTRs which play an important role in post-transcriptional regulation, by controlling translation efficiency(10), subcellular localisation(11), and stability(12). During translation initiation, the 40S ribosome binds mRNA and begins scanning for the first start codon (AUG) and can be interrupted by additional upstream start codons and structures in the 5’UTR(13–15). Thermodynamically stable secondary structures in the UTRs can interrupt the 40S ribosome and decrease translation efficiency depending on their degree of thermodynamic stability(16), length and structure type(17). Addition of the Kozak consensus sequence (GCCGCCACC) surrounding the start codon can also improve ribosome binding and initiation(18).

The m^7^G cap of an mRNA plays essential roles in translation initiation, nuclear export, and stability. The eIF4F complex recognises the cap and recruits the ribosome to the mRNA, making the cap a crucial component of translation initiation(19). The cap also interacts with the cap-binding complex, facilitating mRNA transport from the nucleus into the cytoplasm(20). Finally, the cap protects mRNA from exonuclease-mediated degradation, thereby regulating stability(21). Modification of the mRNA cap structure, particularly using anti-reverse cap analog (ARCA) which is methylated at the 3’ position of the m7G, prevents RNA elongation and ensures that the cap is in the correct orientation. ARCA significantly enhances translation by preventing premature decapping, thereby stabilizing the mRNA transcript(22). The cap1 structure is integral in preventing an innate immune response against foreign RNA(23). ARCA capping initially produces cap0 structure and requires an additional methyltransferase reaction to result in cap1. More recent efforts have improved this cap structure and synthesis, for example, CleanCap cap analogs, which synthesise cap1 in one step. The efficiency of capping has been shown to be more efficient with CleanCap (>95%) compared to ARCA (50-80%), although the extent of benefit on translation efficiency and yield is still controversial(24).

In eukaryotes, the mRNA poly(A) tail is brought into close proximity to the cap, by the cap-binding protein eIF4E and PABPCs, which interact with eIF4G, resulting in the mRNA forming a closed-loop structure in the translation initiation complex(25). The length of poly(A) tail influences mRNA stability and translational efficiency, with longer tails shown to generally enhance both attributes(26, 27). While the consensus used to be that naturally occurring mRNAs have poly(A) tails around 150-250 nucleotides long(28), recently it has been shown that the common length range is 50-100 nucleotides(29), especially those of housekeeping genes(30). These lengths are permissive to the binding of at least one PABC which protects the poly(A) tail from deadenylation(31–34).

The incorporation of modified nucleotides in mRNAs such as pseudouridine (pU), N1- methylpseudouridine (N1mU), 5-methyluridine and 5-methylcytidine (m5C) has been instrumental in driving mRNA therapeutic advancement. When introduced to the cell, unmodified mRNA causes high immunogenicity due to stimulation of Toll-like receptors (TLRs), TLR3, TLR7, and TLR8, ultimately leading to low translational effectiveness. Chemical modifications can dramatically reduce the immunogenicity of mRNA in mammalian cell lines and *in vivo* in mice(35). These chemical modifications can help evade innate immune detection and activation of endosomal TLR3, minimizing inflammatory responses and extending the mRNA’s translational activity through enhanced mRNA stability and protein expression(36). However, over 170 RNA modifications have been identified and understanding their diverse effects on translation is not trivial(37). Recently it was shown that N1mU causes a +1 ribosomal frameshifting at ribosome slippery sequences, likely due to ribosomal stalling(38), highlighting the impact of modified nucleotides on mRNA fidelity and off-target effects and thus the need for sequence optimisation.

Codon optimisation and mRNA structure adjustment play crucial roles in mRNA therapeutic optimisation. By selecting codons that are more frequently used in the target organism, mRNA transcripts can be more efficiently translated(39). Meanwhile, minimizing secondary structures close to the ribosomal binding site through careful sequence modifications can enhance ribosomal access and translation efficiency(40). Maintaining a low overall minimum free energy (MFE) of the transcript with a highly structured coding sequence can stabilise mRNA and prevent premature degradation(41), though this may be at the expense of in-cell protein output(42). Tools such as mRNA Optimiser(43), LinearDesign(9), and mRNAid(8) have allowed streamlined optimisation of codons for a given protein sequence, measured by the codon adaptation index (CAI)(44), and structure for stability, measured by the lowest MFE(45).

Several molecular assays have been adopted to assess therapeutic mRNA translation, however, these assays are often low-throughput and require specialised equipment and computational expertise. Furthermore, these assays lack the ability to produce live recordings of the translation of an mRNA of interest without relying on a reporter protein. Here, we describe the use of HiBit technology to quantify mRNA translation *in vitro* through the rabbit reticulocyte system and *in cellulo* in HEK293 cells constitutively expressing LgBit (Graphical Abstract). HiBit is a component of NanoLuc, an engineered luciferase, that can be introduced as an 11 amino acid tag at either terminus of a POI. In the presence of the LgBit component, and a substrate, HiBit:LgBit dimers form a functional luciferase that emits light in the visible spectra and can be detected using a luminometer(46, 47). The HiBit technology has been extensively used in the study of protein dynamics, especially for measuring protein degradation as brought about by PROTACs(48). The system here described can detect changes in translational and post-translational kinetics of mRNAs encoding different POI and possessing various non-coding elements, making it well suited for high-throughput screening and optimisation of mRNA-based therapeutics.

## Materials and Methods

### Template plasmid construction for linear mRNA

Plasmids (pBluescript KS+, pBS hereon) for HiBit-tagged POI mRNA were generated by Gibson Assembly (Gibson Assembly cloning kit, NEB, E5510S) or restriction digest cloning. Fragments consisting of digested pBS backbone, PCR product 5’UTR-POI-spacer-HiBit and PCR product HiBit-3’UTR-AgeI were assembled by Gibson assembly according to the manufacturer’s instructions. Specifically, POI-spacer-HiBit fragments were amplified from previously cloned POI-HiBit fragments in pcDNA3 backbone. The 3’UTR fragment was amplified from previously generated pBS-5’UTR_eGFP_3’UTR plasmid(49). All PCRs were performed with Q5 High-Fidelity DNA Polymerase (NEB, M0491S) and digested with DpnI (NEB, R0176S). PCR and digest products were purified using Monarch DNA Gel Extraction and PCR and DNA Clean Up Kits (NEB T1020S, T1030S).

To generate the pBS-eGFP-HiBit-3’UTR construct lacking a 5’UTR (No 5’UTR), a PCR was performed using a forward primer lacking the 5’UTR but with the Kozak upstream of the ATG. The PCR product was cloned into the pBS backbone by restriction digest and ligation using T4 DNA ligase (NEB, M0202S). For the construct with a structured 5’UTR (5’UTR-Hairpin), RNAfold(50) webserver was used to predict the hairpin structure. The design consisted of a sequence forming a hairpin containing the Kozak sequence in the loop with the start codon immediately downstream within the stem region of the hairpin. The fragment was synthesised by Thermo Fisher Gene Strings and cloned into the pBS backbone by restriction enzyme cloning.

For MycT58A mRNA optimisation, the LinearDesign algorithm(9) was run with the MycT58A protein sequence using Python (version 2.7) and Clang (version 17.0.6). For CAI optimisation a Lambda value of 100 was chosen, and for MFE optimisation a value of 0 was chosen. Myc T58A CAI and MFE were manufactured by GENEWIZ as gene synthesis constructs and cloned into pBS using restriction digest cloning. All constructs generated were confirmed by Sanger sequencing.

### Synthesis and purification of linear mRNA

For mRNA synthesis, 1 µg of plasmids pBS-5’UTR-POI-HiBit-3’UTR were linearised by restriction enzyme digest and purified using QIAGEN QIAquick PCR Purification Kit (Qiagen, 28104) according to the manufacturer’s instructions. From the linearised plasmid, the template for mRNA synthesis was amplified by PCR with Platinum Taq DNA Polymerase High Fidelity (Invitrogen, 11304-011) according to the manufacturer’s programme specifications and the following primers.

T7-Fwd: 5’-GTGAGCGCGCGTAATACG-3’ PolyT-3T-Rev: 5’-TTTTCCCTCACTTCCTACTCAGGC-3’ PolyT-40T-Rev: 5’- TTTTTTTTTTTTTTTTTTTTTTTTTTTTTTTTTTTTTTTTTCCCTCACTTCCTACTCAGGC-3’ PolyT-80T-Rev: 5’- TTTTTTTTTTTTTTTTTTTTTTTTTTTTTTTTTTTTTTTTTTTTTTTTTTTTTTTTTTTTTTTTTTTTTTTTTTTTTTTTTCCCTCACTTCCTACTCAGGC-3’ PolyT-120T-Rev: 5’- TTTTTTTTTTTTTTTTTTTTTTTTTTTTTTTTTTTTTTTTTTTTTTTTTTTTTTTTTTTTTTTTTTTTTTTTTTTTTTTTTTTTTTTTTTTTTTTTTTTTTTTTTTTTTTTTTTTTTTTTTCCCTCACTTCCTACTCAGGC-3’

Following PCR clean-up, MEGAscript T7 Transcription Kit (Invitrogen, AM1333) was used to perform *in vitro* transcription for mRNA synthesis (49), with an incubation of 16-18h. Several components of the mRNA synthesis were used in combination in this reaction: ARCA (NEB, S1411), CleanCap Reagent AG (Trilink, N-7113), Pseudouridine-5’-Triphosphate (Trilink, N- 1019), N1-Methylpseudouridine-5’-Triphosphate (Trilink, N-1081), 5-Methoxyuridine-5’- Triphosphate (Trilink, N-1093), 5-Methylcytidine-5’-Triphosphate (Trilink, N-1014). Following the *in vitro* transcription, samples were treated with TURBO DNase (ThermoFisher, AM2238) for 15 minutes to degrade template DNA from the synthesis product. mRNA was purified using MEGAclear Transcription Clean-Up Kit (Invitrogen, AM1908) following the manufacturer’s instructions. Purified mRNA was treated with Antarctic Phosphatase (NEB, M0289) for 1h at 37°C followed by another purification round using MEGAclear. mRNA concentration was determined using a NanoDrop spectrophotometer (Thermo Scientific). Correct mRNA size was confirmed by electrophoresis by running on 1% non-denaturing agarose gel. Purified mRNA was stored at -80°C.

### *In vitro* HiBit translation assay in Rabbit Reticulocyte Lysates (RRL)

The *in vitro* translation assay was performed by adapting the protocol of Promega’s Flexi Rabbit reticulocyte lysate kit (nuclease treated, L4540). Given the different sizes of the mRNA products, the comparison between constructs must be made with equal molecules of mRNA per reaction, for this we recommend using 1.5pmol of mRNA per reaction. First, mRNA and all components of the kit were thawed on ice. A reaction master mix was prepared and kept on ice as detailed in table 1. For time course experiments, we recommend preparing the master mix to include as many mRNA samples as time points, plus one for pipetting errors, given the viscosity of the reticulocyte lysate mix.

**Table 1.** Reagents and volume of components of a RRL reaction.

| Component | mRNA sample (μL) | No RNA control (μL) |
| --- | --- | --- |
| Flexi Rabbit Reticulocyte Lysate | 8.25 | 8.25 |
| Amino acid mixture Minus Leucine<br>(1 mM) | 0.25 | 0.25 |
| Amino acid mixture Minus Methionine<br>(1 mM) | 0.25 | 0.25 |
| KCl (2.5 M) | 0.5 | 0.5 |
| Ribonuclease inhibitor (Thermo<br>Fisher Scientific, N8080119) | 0.5 | 0.5 |
| MgACO (25 mM) | 0.5 | 0.5 |
| DTT (100 mM) | 0.25 | 0.25 |
| RNA (0.75 pmol/μL) | 2 | 0 |
| Nuclease-Free H<sub>2</sub>O | 0 | 2 |
| **Final V** | **12.5** | **12.5** |

Once prepared, the master mix was divided into separate tubes for each time point or mRNA condition, all kept on ice. For experiments aimed at comparing the translational efficiency of different mRNAs at a single time point, a master mix lacking mRNA was prepared and divided into separate tubes (10.5 µL/tube) to which 2 µL of mRNA were added separately to each tube. All tubes were then added to a heat block set at 30°C for the duration of the incubation and immediately placed back on ice. For time course experiments, appropriate tubes were removed according to the equivalent time point. A single time point of No RNA control with incubation of 120 minutes was run. Once all samples had cooled, they were stored at -80°C to ensure termination of the reaction. Translation reaction replicates were performed on different days, with each sample run in duplicate, and measured on a plate-based luminometer. On ice, the resulting HiBit protein was measured by placing 2.5 µL of each sample in a well of a white-walled 96-well plate (Nunc™ MicroWell™ 96-Well, Nunclon Delta-Treated, Flat-Bottom Microplate, 136101) in duplicate or triplicate. To minimise variability all experimental replicates were processed in the same plate. To all wells, 50 µL of HiBit buffer (LgBit 1:200, Promega, N112A; Furimazine 1:100 Promega, N113A in nuclease-free water) were added. Samples and buffer were mixed briefly on a shaker maintaining samples on ice, then luminescence readings were acquired using a CLARIOstar Plus luminometer (530–540).

### Western blotting

For Western blotting of RRL samples, 1.6 µL of each replicate lysate were pooled to a final 5 µL volume, Sample Buffer (50 mM Tris-HCl pH6.8, 2% SDS, 0.1% bromophenol blue, 10% glycerol, 100mM dithiothreitol added fresh) was added to bring to a final volume of 25 µL, then samples were heated to 80°C for 2 minutes for protein denaturation. Of denatured samples, 4 µL were loaded on a 10-15% acrylamide/bis gel (for better resolution of eGFP and eGFP-HiBit sizes, 15% polyacrylamide gel was used), and proteins were separated by electrophoresis at 150V for 1.5-3h, depending on the gel’s acrylamide percentage. Proteins were transferred to a pre-activated PVDF membrane in 1-Step Transfer buffer (ThermoFisher, 84731) using a semi-dry Pierce G2 Fast Blotter (ThermoFisher), which allows transfer at 25V, 1.3A in 15 minutes according to the manufacturer’s instructions. After transfer, membranes were blocked in 2% BSA in TBS with 0.1% Tween 20 for 1 hour at room temperature and incubated overnight in rabbit anti-eGFP (1:1000, abcam, ab290) and mouse anti-GAPDH (1:5000, Proteintech, 60004-1-Ig). Following incubation with primary antibodies, membranes were washed with TBS with 0.1% Tween 20 three times, with each wash lasting 5 minutes, followed by 1 hour incubation with secondary antibody solution (0.01% SDS, 20% FBS, TBS with 0.1% Tween 20) containing fluorescent secondary antibodies IRDye 680RD Goat anti-Rabbit IgG Secondary Antibody (1:20 000, Li-COR Biosciences 926-68071) and IRDye 800CW Goat anti-Mouse IgG Secondary Antibody (1:10 000, Li-COR Biosciences 926-32210). Following incubation with secondary antibodies, membranes were washed with TBS with 0.1% Tween 20 three times and imaged using an Odyssey Li-COR Imaging System (Li-COR Biosciences). Following imaging of the fluorescent signal, membranes were incubated for 1 hour with LgBit buffer (0.5% LgBit in TBS with 0.1% Tween 20). In the last 5 minutes of incubation in LgBit buffer, Furimazine substrate was added to the buffer at a 500-fold dilution. The membranes were imaged again for bioluminescence recording of HiBit:LgBit signal using the same imaging system. The resulting images were analysed on ImageStudioLite (Li-COR Biosciences) by measuring the average fluorescence intensity of a region of interest (ROI) around the respective band and subtracting the intensity reading of an ROI of the same size in the No RNA control. The background-corrected signal was then normalised to GAPDH loading control.

### Live *in cellulo* HiBit translation assays

HEK293 LgBit cells (gift from Itzhaki lab) were cultured in Dulbecco’s Modified Eagle’s Medium (DMEM, ThermoFisher, 41966052) supplemented with L-glutamine (Thermo Fisher, 25030-024) and 10% FBS (Sigma, F7524). For live cell experiments of mRNA translation kinetics, HEK293 LgBit cells were reverse-transfected with 1.2 pmol of mRNA with Lipofectamine RNAiMAX Transfection Reagent according to the manufacturer’s instructions and plated on clear bottom, white-walled plates (Greiner Bio-One CELLSTAR 96-well, White, Cell Culture-Treated, Flat-Bottom Microplate, 655098) at 40, 000 cells/well in Leibovitz’s L-15 Medium no phenol red (ThermoFisher, 21083-027), which supports cell growth in environments without CO2 equilibration, supplemented with 10% FBS and substrate Nano-Glo Endurazine Live Cell Substrate (Promega, N2570). Immediately post-transfection, cells were placed in a CLARIOstar Plus luminometer set to 37°C where recordings were taken every 12 minutes for 18 hours (530–540). Negative controls of mock transfections lacking RNA were performed.

### Quantitative reverse transcription PCR

Total RNA was isolated using NucleoSpin RNA Mini kit (Macherey-Nagel, 74095550) according to the manufacturer’s instructions. Once isolated, NanoDrop Lite spectrophotometer (ThermoFisher Scientific) was used to quantify and assess the purity of RNA. RNA was reverse transcribed using the High Capacity cDNA Reverse Transcription kit (random primers) (ThermoFisher Scientific, 4368814) according to the manufacturer’s instructions. The RT-PCR reactions were performed by the CFX Opus 384 (Bio-Rad) using 5 µL Fast SYBR Green Master Mix (Applied Biosystems, 4385612), 0.5 µL of each oligonucleotide primer, and 1 µL cDNA to a final volume of 10 µL. The 2^-ddCT^ method was used to determine gene expression changes using GAPDH as a housekeeper gene. Primers for target genes (listed below) were designed and synthesised by Merck and were used at a final concentration of 250nM.

GAPDH-Fwd: 5’- GGAGCGAGATCCCTCCAAAAT -3’

GAPDH-Rev: 5’- GGCTGTTGTCATACTTCTCATGG -3’

IFNB1-Fwd: 5’- GCAGTTCCAGAAGGAGGACG -3’

IFNB1-Rev: 5’- AGCCAGGAGGTTCTCAACAAT -3’

RIGI-Fwd: 5’- CTCTTTGTGAAAACCAGAGCACTT -3’

RIGI-Rev: 5’- CGGGAGGGTCATTCCTGTGTT -3’

TL3-Fwd: 5’- TGGCTTGTCATCTACAAAATTAGG -3’

TL3-Rev: 5’- AACACCCTGGAGAAAACTCTTTA -3’

### Statistics

Statistical analyses were performed on normalised data using GraphPad Prism v10.1.1 (GraphPad Software, Inc., San Diego, CA, USA) as indicated with P ≤ 0.05 considered statistically significant.

For *in vitro* HiBit translation experiments, luminescence recordings for each sample were normalised to the average of the respective No RNA control. Given the novelty of this assay, the Z’ (or Z-factor), a statistical measurement of the robustness of assays(51), was calculated by comparing the signals from the minimal and maximal signals recorded in the assays. For the RRL HiBit assay, No RNA and eGFP-HiBit 40A samples respectively were used as minimum and maximal signal, resulting in a Z’=0.8, indicating an optimal dynamic range.

For live, *in cellulo* HiBit translational assays, each condition’s signal was normalised to its first time point as the signal at the beginning is background level for all conditions, to generate fold changes in RLU. Once more, the Z’ was calculated, using the peak and end values (4-hour and 18-hour time points, as plotted in Figure 5C) for eGFP-HiBit with poly(A) of 40A against No RNA control, which resulted in a Z’=0.8.

Statistical analysis of *in cellulo* data normalised to time 0 was performed at 4 hours or 18 hours by One-way analysis of variance (ANOVA) with multiple comparisons made to the reference HiBit mRNA. Furthermore, to assess the contribution of coding and non-coding elements to the translation of POI-HiBit at 4 hours and 18 hours, relative luminescence units were normalised to the average of the reference construct and followed by Two-way ANOVA with matched values, to reflect continuous data.

## Results

### The addition of HiBit does not hinder the translation of eGFP *in vitro* and offers a high-throughput system for monitoring translational kinetics

To determine whether the addition of a tag encoding HiBit at the 3’ end of eGFP-NLS alters the translation kinetics of the fluorophore, two mRNA expressing eGFP and eGFP-HiBit were synthesised. These mRNAs possess identical 5’- and 3’-UTR and poly(A) tail lengths (120 adenosines) and differ only in the presence of a serine and glycine linker and the HiBit coding sequence. We employed the cell-free *in vitro* translation assay RRL system to evaluate the rate of translation of the mRNAs over a time course (1, 5, 10, 20, 30, 60, 90 and 120 minutes). Western blotting antibody-based detection of proteins was performed, followed by direct, incubation with LgBit and substrate Furimazine to detect HiBit-tagged eGFP by bioluminescence (Figure 1A). Both methods provided an accurate and sensitive measurement of HiBit-tagged protein, however, direct detection of eGFP-HiBit does not rely on an antibody against a POI and confirmed that the higher molecular weight band detected by the GFP antibody was eGFP-HiBit. Quantification of the efficiency of translation confirmed that the addition of HiBit did not affect the translation kinetics of eGFP (Figure 1B).

**Figure 1:**
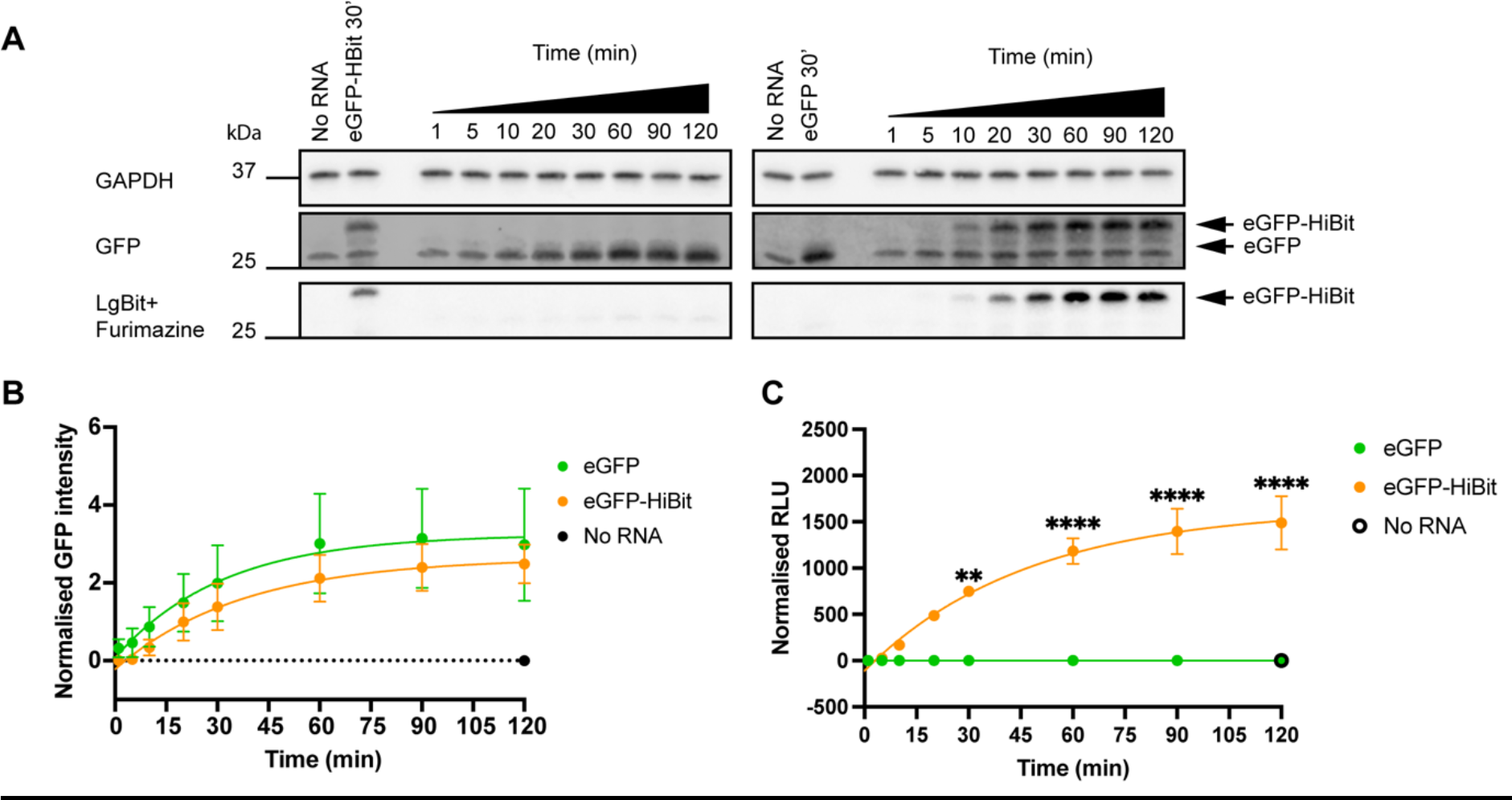
The addition of the HiBit tag allows the detection of POI-specific translational kinetics profiles. (A) Translation kinetics of eGFP and eGFP-HiBit in the RRL system by western blotting and quantified in **(B)** normalised to GAPDH, eGFP (green) or eGFP-HiBit (orange). (**C**) Translation kinetics of mRNA encoding eGFP (green) or eGFP-HiBit (orange) measured by luminometer, Relative Luminescence Units (RLU) of all samples were normalised to No RNA sample. Mean and Standard Error of the Mean (SEM) of three biological replicates are plotted and Two-way ANOVA was conducted to compare eGFP and eGFP-HiBit signal at each time point. ** p<0.005, **** p<0.0001.

Despite the method sensitivity, Western blotting results in high variability (large error bars), is low-throughput and time-consuming, therefore, a plate-based assay was developed. For this, RRL samples were incubated with LgBit and substrate Furimazine, for 5 minutes before luminescence readings were acquired. The assay produced less variable results and reproduced similar translation kinetics seen by Western blotting (Figure 1C). The assay specifically detected the HiBit:LgBit-induced luminescence as eGFP samples lacking the HiBit tag did not produce a bioluminescence reading. Importantly, since the assay is compatible with a multi-well format, it allows for triplicate readings, and testing of multiple experimental replicates or samples in one plate, making this assay optimal for high-throughput work.

### HiBit-encoding mRNA can be used to study the contribution of the non-coding and coding elements of mRNA to its translation *in vitro* and *in cellulo*

#### CAP and 5’ UTR

To further validate the sensitivity of the HiBit assay, we evaluated the contribution of cap-dependent translation by comparing eGFP-HiBit mRNA with no cap, ARCA (anti-reverse cap analog), and CleanCap AG in the RRL system and measured on a plate-based luminometer (Figure 2A-C). We determined the cap is crucial for efficient protein translation, and the absence of cap significantly (p<0.0001) reduced the detected luminescent signal (Figure 2A, B) and protein level (Figure 2C). However, no significant difference was measured between ARCA and CleanCap in this *in vitro* system (Figure 2B).

**Figure 2:**
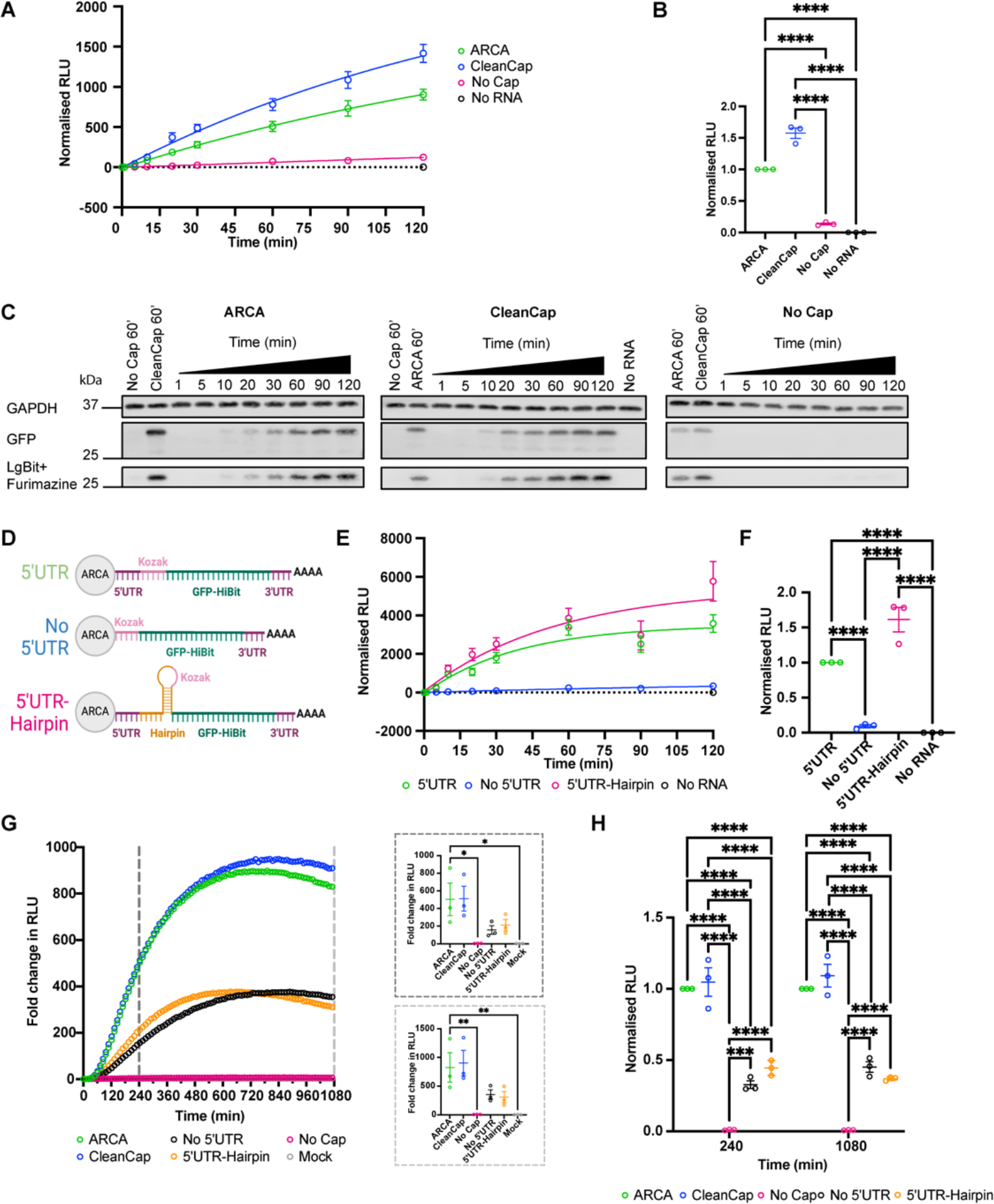
HiBit-encoding mRNA can detect alterations to the 5’UTR. (A) Translation kinetics of mRNA capped using ARCA (green), CleanCap (blue) or uncapped mRNA (magenta) encoding eGFP-HiBit were compared in the RRL system; RLU was normalised to No RNA control. **(B)** Scatter plot depicting RLU of three biological replicates at 120 minutes normalised to ARCA sample, Mean and SEM of three biological replicates are plotted and Ordinary one-way ANOVA was conducted. **(C)** Western blot of pooled RRL samples with GAPDH as a loading control, with eGFP-HiBit protein detected with an anti-GFP antibody and LgBit and Furimazine. **(D)** Schematic representation of the mRNA structures with a 5’UTR, No 5’UTR and 5’UTR-Hairpin. **(E)** mRNA translation kinetics of 5’UTR, no 5’UTR and 5’UTR-Hairpin constructs were compared in the RRL. RLU was normalised to No RNA control. **(F)** Scatter plot depicting RLU of three biological replicates at 120 minutes normalised to 5’UTR mRNA, Mean and SEM, and Ordinary one-way ANOVA. **(G)** Fold change in RLU from time 0 of HEK293 LgBit cells transfected with eGFP-HiBit mRNA with: ARCA-capped (green), CleanCap-capped (blue), no Cap (Magenta), no 5’UTR (black) or 5’UTR-Hairpin (orange) or mock (grey). Mean fold change RLU per time point of three biological replicates are shown for the complete time course; Mean and SEM are depicted at 4 hours (dark grey dashed line) and 18 hours (light grey dashed line), where Ordinary one-way ANOVA compared to reference sample (ARCA) was conducted. **(H)** Mean and SEM of RLU normalised to ARCA sample at 240 and 1080 minutes were compared by Two-way ANOVA with matched values. * p<0.05, ** p<0.005*** 0.0001<p<0.0005, **** P<0.0001.

To further challenge the HiBit assay system, the properties of the 5’UTR were altered by designing two new mRNAs, one without a 5’UTR (No 5’UTR) and one which contained an 18bp complementary region resulting in a hairpin structure (5’UTR-Hairpin) engineered immediately adjacent to the 5’UTR, around the start codon (Figure 2D). The thermal stability and free energy of these sequences were estimated using the RNAfold(50). The 5’UTR is predicted to have an MFE of -7.7 kcal/mol and thermodynamic ensemble free energy of -9.04 kcal/mol. The 5’UTR-Hairpin is predicted to have an MFE of -43.4 kcal/mol and thermodynamic ensemble free energy of -46.89 kcal/mol. The translational kinetics of these constructs were measured with the HiBit RRL system. No HiBit:LgBit signal could be detected in the absence of the 5’UTR, indicating little to no protein was synthesised over the course of the experiment, while both mRNAs with intact 5’UTRs generated a similar translation profile (Figure 2E). When comparing the luminescent signal at 120 minutes, no significant difference was detected between mRNA with 5’UTR and 5’UTR-Hairpin, but both constructs produced significantly higher signal than No 5’UTR and No RNA control (p<0.0001, Figure 2F).

To detect translation in live cells, we harnessed the HiBit system to express HiBit-tagged proteins in HEK293 cells constitutively expressing LgBit. The experimental method allows the live recording of the kinetics of translation and post-translational processing of the transfected mRNA and resulting HiBit-tagged protein. HEK293 LgBit cells were reverse-transfected with the eGFP-HiBit mRNAs (ARCA, CleanCap, No Cap, No 5’UTR and 5’UTR-Hairpin) and treated with a stable substrate for the luciferase dimer, Endurazine. The mRNA with No Cap significantly abolished translation (p<0.0001), whilst mRNA capped with ARCA and CleanCap resulted in a rapid increase in protein production, reaching plateau at 720 minutes (Figure 2G- H). Contrary to the RRL, the presence of a structured element adjacent to the 5’UTR (5’UTR- Hairpin) in addition to the absence of the 5’UTR (No 5’UTR) resulted in significantly slower translational kinetics compared to ARCA-capped mRNA with the 5’UTR (Figure 2G, H). Specifically, removing the 5’UTR resulted in a significant reduction in total protein yield luminescence (p<0.0001), with a 67% decrease at 4 hours during the exponential translation phase, and 55% at 18 hours. Introducing a hairpin decreased signal by 56% and 63% at 4h and 18h, respectively (Figure 2H). These results highlight the ability of this novel sensitive cellular assay to measure changes in the translation efficiency of mRNAs with differing non-coding elements at the 5’ end.

#### Coding sequence

Given that eGFP is highly engineered and its translation is extremely efficient (ARCA and CleanCap constructs, Figure 2G), an mRNA encoding a non-codon-optimised, much less stable protein, human oncogene cMyc (Myc), was generated to investigate translation kinetics and efficiency. All constructs retained the identical Cap (ARCA), 5’UTR, linker and HiBit sequence and, 3’UTR as described for eGFP-HiBit. eGFP-HiBit and Myc-HiBit encoding mRNAs were translated for 120 minutes in the RRL system and measured on a plate-based luminometer. The efficiency of eGFP-HiBit translation was far greater than that of Myc-HiBit (Figure 3A, p=0.0002). In HEK293 LgBit cells, eGFP-HiBit encoding mRNA resulted in a rapidly translated protein that quickly reached a plateau when protein synthesis and degradation reached a steady state (Figure 3B). In agreement with the observation within the RRLs, translation of Myc-HiBit mRNA resulted in a significantly lower (p<0.0001) luminescent signal in cells both during the exponential translation phase (4 hours, Figure 3B dark grey call- out) and at the end of the experiment (Figure 3B light grey call-out).

**Figure 3:**
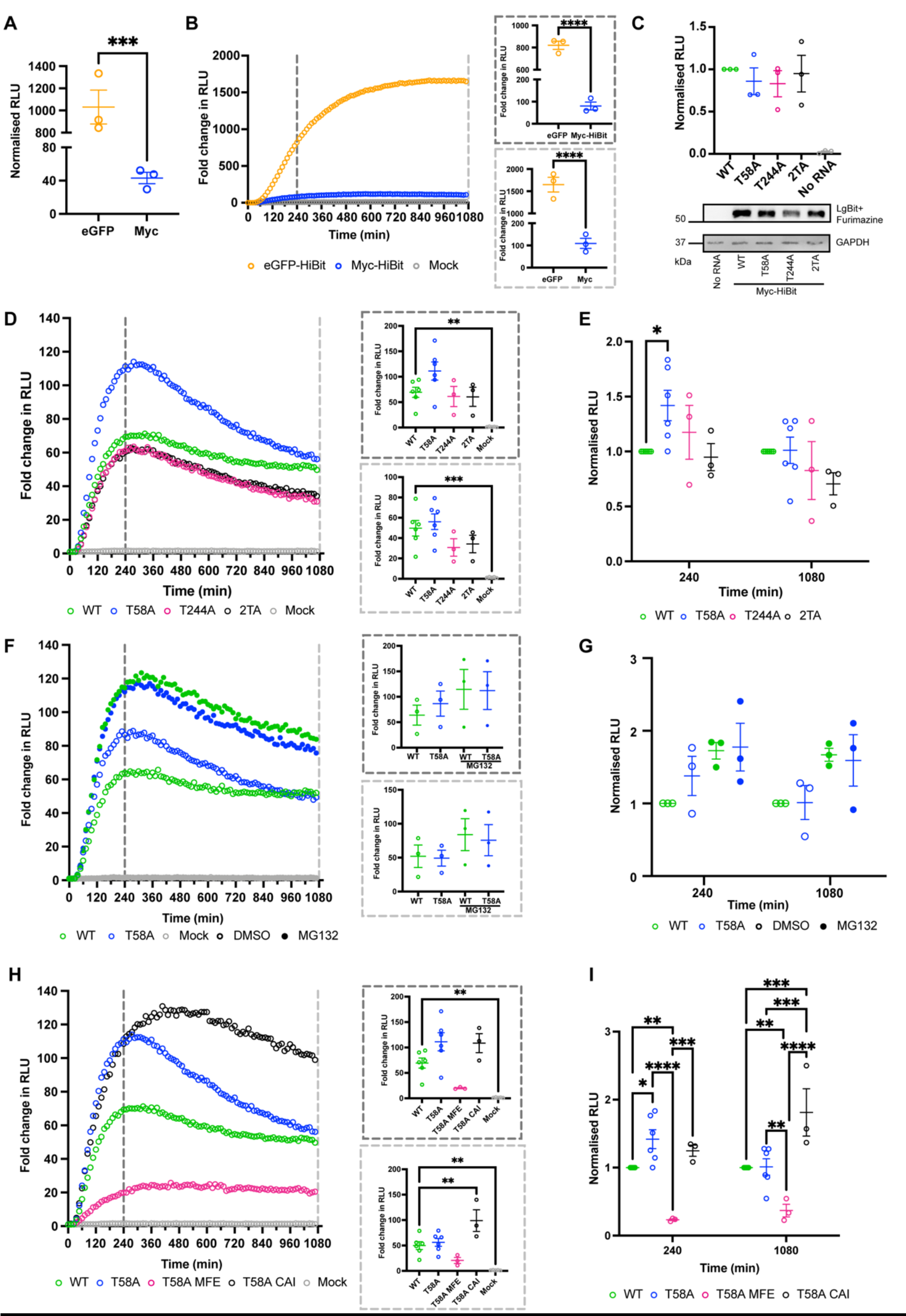
HiBit-encoding mRNA reports on optimisation of Myc coding sequence. (A) RRL reactions were carried out for 120 minutes with mRNA encoding eGFP-HiBit and Myc- HiBit, RLU was normalised to No RNA; scatter plots with SEM bars represent the mean from three independent biological replicates. **(B)** Fold change in RLU from time 0 of HEK293 LgBit cells transfected with eGFP-HiBit and Myc-HiBit. Mean and SEM per time point of three biological replicates are shown for the complete time course and at 4 hours (dark grey dashed line) and 18 hours (light grey dashed line), where Ordinary one-way ANOVA was conducted. **(C)** mRNA translation of Myc with threonine (T) to alanine (A) substitutions at key phosphodegron sites were compared in the RRL system by incubating the HiBit mRNA for 120 minutes. Scatter plot depicting RLU of three biological replicates normalised to WT Myc-HiBit, mean and SEM. Western blot of pooled samples from the three replicates, displaying GAPDH and Myc-HiBit detected by LgBit and Furimazine. **(D)** Live cellular kinetic assay in HEK293 LgBit cells transfected with Myc-HiBit or Mock, data normalised to t0; dots representing the mean of at least three biological replicates per time point. Mean and SEM at 4 hours (dark grey dashed line) and 18 hours (light grey dashed line) are shown and analysed by ordinary One-way ANOVA compared to WT Myc-HiBit. **(E)** Data at 4 hours and 8 hours was normalised to WT to depict changes in Myc-HiBit signal and compared using Two-way ANOVA with matched values. **(F)** HEK293 LgBit transfected with Myc-HiBit WT or T58A were co-treated with DMSO (0.005%, empty circles) or MG132 (Cf= 4µM, full circles) at seeding and expression was measured through time and normalised to t0 and compared to WT by One- way ANOVA. **(G)** Data at 4 hours and 8 hours was normalised to WT DMSO and compared by Two-way ANOVA with matched values. **(H)** Two codon optimisation strategies applied to Myc T58A and translation were compared in the live cellular kinetic assay in HEK293 LgBit cells. Fold change in RLU from time 0 is shown with dots representing the mean of three biological replicates per time point. Mean and SEM at 4 hours (dark grey dashed line) and 18 hours (light grey dashed line) are shown and compared to reference WT by One-way ANOVA. **(I)** Data at 4 hours and 8 hours was normalised to WT to depict changes in Myc-HiBit levels and compared using Two-way ANOVA with matched values. * p<0.05, ** p<0.005. *** 0.0001<p<0.0005, **** p<0.0001. WT = Wild-type Myc (green); T58A = Threonine-to-Alanine substitution at T58 of Myc (blue), T244A = Threonine-to-Alanine substitution at T244 (magenta in D, E); 2TA = Threonine-to-Alanine substitution at T58 and T244 (black in D, E); T58A MFE = Minimum-free-energy optimisation of T58A Myc (magenta in H, I), T58A CAI = Codon- adaptation-index optimisation of T58A Myc (black in H, I).

We next investigated the HiBit assay’s sensitivity to subtle substitutions in coding sequences. We focused on the Myc coding sequence since eGFP is optimised for translation and protein stability. The rapid protein turnover of Myc is, in part, mediated by the phosphodegron site threonine (T) 58, which is a recognition site for E3 ligase, FBW7(52–54). More recently a second phosphodegron for regulation of Myc by FBW7 has been discovered(55) at threonine 244. To determine the effects of phosphoablation on these phosphodegron sites of Myc-HiBit, we generated 3 Myc mutants – T58A, T244A and 2TA (T58A and T244A). Using the RRL system, no significant difference was recorded when comparing mutants to WT Myc (Figure 3C), suggesting all mRNAs are equally translated. Myc-HiBit mRNAs were then tested in the live kinetic cellular assay. All versions of Myc were rapidly translated with peak signal reached around 4 hours (Figure 3D), However, the signal rapidly declined suggesting that the longevity of the protein signal may depend on the protein half-life of the encoded POI. Interestingly, none of the stabilising mutations of Myc-HiBit resulted in an improved, more sustained signal at 18h however, T58A alone surpassed WT Myc expression at the 4-hour peak (Figure 3D). When comparing the mutants to WT at these time points (Figure 3E), T58A resulted in a significantly higher (p=0.0474) average of 42% increase in stability at 4 hours which is reduced to a negligible 1% at 18 hours. Myc T244A only resulted in a 17.5% increase in average signal at 4 hours and an 18% loss of average signal at 18 hours. The double phosphodegron mutant 2TA had comparable values to WT at 4 hours (average 5% less than WT) but dropped steeply to 30% less average signal at 18 hours.

To prove that the observed difference between Myc WT and T58A is due to the changes in post-translational regulation, HEK293 LgBit cells were treated with a sublethal dose of proteasome inhibitor MG132 (4 µM) or vehicle control DMSO at the time of transfection. Proteasomal inhibition was demonstrated by the resulting increased WT Myc-HiBit signal, elevated by 73% at 4 hours and 67% at 18 hours. As predicted, at 4 hours, there is an overlap in WT and T58A signal upon MG132 treatment, indicating that the higher peak of T58A in the DMSO control is due to reduced degradation of the mutant compared to the WT at comparable exponential translation rates (Figure 3F-G).

Since regulatory elements have been reported to control Myc mRNA turnover(54, 56–59), Myc-HiBit mRNA optimisation strategies were shifted to improving mRNA translation and stability. Two mRNA optimisation strategies, aimed at improving mRNA stability and codon optimality(9), were applied to the best stability mutant, Myc T58A-HiBit. Applying the published algorithm, we generated mRNA encoding for the sequence with the highest codon adaptability index (CAI) and the highest minimum-free-energy change (MFE). Specifically, the MFE and CAI indexes respectively were: -509.00 kcal/mol; 1.000 for T58A CAI; -944.40 kcal/mol; 0.737 for T58A MFE. Importantly, only the open reading frame of Myc T58A sequence was optimised (-457.60 kcal/mol; 0.819 MFE and CAI indexes), whilst the UTRs, poly(A), linker and HiBit sequence remained unchanged to allow for accurate comparison to WT and T58A Myc-HiBit mRNA. Optimisation of the mRNA stability with T58A MFE impeded efficient translation (Figure 3H), significantly reducing the expression of Myc-HiBit by 77% compared to WT at 4h (p=0.0012), and 63% at 18h (p=0.0083, Figure 3I). However, despite the reduced protein abundance, over the time course of the experiment, there was no rapid decline in signal suggesting an enhanced mRNA stability. Improving the codon adaptability index (T58A CAI) resulted in a delayed peak at 8 hours compared to WT and T58A Myc-HiBit and a more sustained signal over time (Figure 3H). Specifically, at 4h, T58A CAI showed an average 25% increase in luminescence signal over WT compared to the 42% of T58A (Figure 3I), but at 8h there is a statistically significant (p<0.0001) 57% increase in signal over WT with T58A CAI against the 30% of T58A. At 18h, there is a statistically significant improvement of T58A CAI over all versions of Myc-HiBit, with an average 81% increase in WT Myc-HiBit signal (p=0.006) which is not seen with non-codon optimised T58A.

#### Modified nucleotides

The impact of nucleoside modifications on the expression of the encoded protein is likely dependent on the gene of interest. Employing the HiBit system, full substitutions of unmodified uridine or cytidine with modified analogues were implemented during the *in vitro* transcription reaction of eGFP-HiBit and Myc-HiBit. Specifically, the contribution of Pseudouridine-5’- Triphosphate (pU), N1-Methylpseudouridine-5’-Triphosphate (N1mU), 5-Methoxyuridine-5’- Triphosphate (5moU) and 5-Methylcytidine-5’-Triphosphate (m5C) on translation were investigated in the HiBit assay system. Unmodified (Un) or modified RNAs (modRNA) were translated *in vitro* with the RRL system for 120 minutes and resulting HiBit-tagged proteins were detected via a luminometer (Figure 4A) and by Western blotting (Figure 4B). All modifications to eGFP-HiBit and Myc-HiBit resulted in a translated protein. For both mRNAs, modification with m5C resulted in a non-significant reduction in translated protein, with 37% and 57% respective average decrease in luminescent signal (Figure 4A). This trend was confirmed by Western blotting of the pooled samples (Figure 4B).

**Figure 4:**
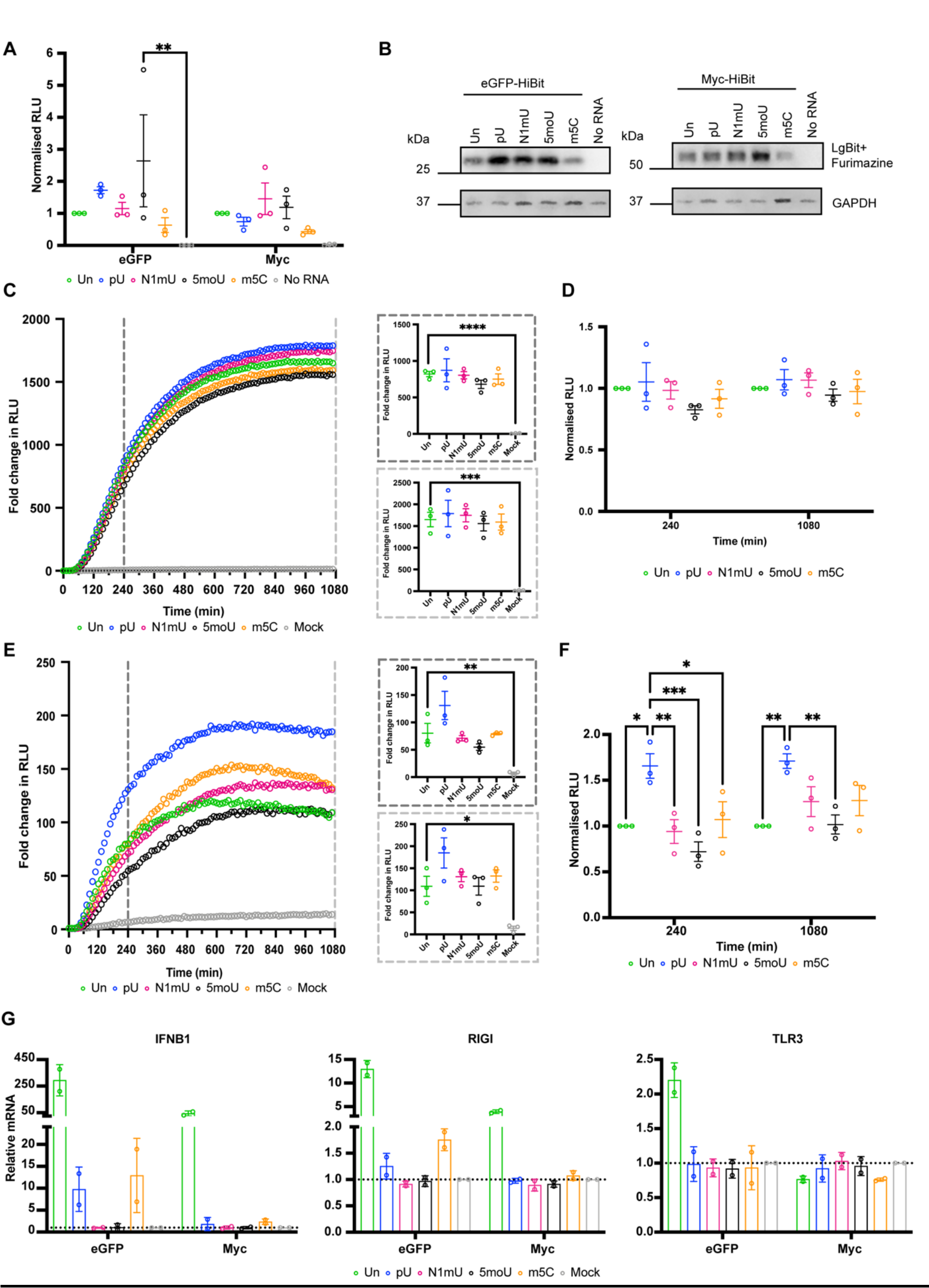
Effect of nucleotide modifications on HiBit-encoding mRNA translation depends on the encoded POI. **(A)** mRNA translation of eGFP-HiBit or Myc-HiBit mRNA containing unmodified or 100% substitution with modified nucleotides was compared in the RRL system by incubating the HiBit mRNA for 120 minutes. Scatter plot depicting RLU of three biological replicates normalised to Unmodified (Un) eGFP-HiBIt or Myc-HiBit, mean and SEM. **(B)** Western blot of pooled samples from three RRL replicates showing GAPDH and eGFP- HiBit or Myc-HiBit detected by LgBit and Furimazine. **(C)** Live cellular kinetic assay in HEK293 LgBit cells transfected with eGFP-HiBit (unmodified and modified) or Mock, data normalised to t0; dots representing the mean of three biological replicates per time point. Mean and SEM at 4 hours (dark grey dashed line) and 18 hours (light grey dashed line) are shown, ordinary One-way ANOVA compared to Unmodified eGFP-HiBit. **(D)** Data at 4 hours and 8 hours was normalised to Unmodified to depict changes in eGFP-HiBit signal and compared by Two-way ANOVA with matched values. **(E)** HEK293 LgBit transfected with Myc-HiBit (unmodified or modified) or Mock and fold change in RLU from t0 was plotted and compared to WT by Ordinary One-way ANOVA. **(F)** Data at 4 hours and 8 hours was normalised to unmodified and compared using Two-way ANOVA with matched values. **(G)** Induction of immunogenic gene expression (IFNB1, RIGI and TLR3) in HEK1293 LgBit cells transfected with unmodified or modified eGFP-HiBit or Myc-HiBit was assessed by qPCR and normalised against mock sample. * p<0.05; ** p<0.005; *** 0.0001<p<0.0005; **** p<0.0001. Un= Unmodified (green); pU = Pseudouridine-5’-Triphosphate (blue); N1mU = N1-Methylpseudouridine-5’-Triphosphate (magenta); 5moU = 5-Methoxyuridine-5’-Triphosphate (black); m5c = 5-Methylcytidine-5’- Triphosphate (orange).

Next, we assessed the effect of the modifications on mRNA translation kinetics *in cellulo*. Incorporation of modifications into eGFP-HiBit did not affect its translational kinetics profile, which displayed rapid accumulation and plateauing. All modifications showed (Figure 4C) no significant changes compared with unmodified mRNA (Figure 4D). When comparing the effects of nucleotides on Myc-HiBit mRNA expression, a greater distinction in the effect of the modifications was observed (Figures 4E-F). Myc-HiBit expression was recorded for all mRNAs regardless of modification status, however, pU substitution produced both a faster initial burst of Myc-HiBit synthesis and a more prolonged steady-state of synthesis over the time course of the assay (Figure 4E), resulting in an increased average Myc-HiBit signal of 66% at 4 hours (p=0.0103), and further 71% at 18 hours (p=0.0052) when compared to unmodified Myc-HiBit (Figure 4F). Since improved translation rates do not solely dictate the choice of modification, we verified the immunogenic response to all mRNAs by qRT-PCR. We selected three genes which encode for cytosolic RNA sensors (retinoic acid-inducible gene I, RIGI; TLR3) and downstream effectors (interferon beta 1, IFNB1). mRNAs encoded with unmodified nucleotides resulted in a large immunogenic response, which was substantially higher for eGFP-HiBit than Myc-HiBit modRNA (Figure 4G). The inclusion of modified nucleotides blunted the immunogenic response to varying degrees dependent on modification (Figure 4G).

#### Poly(A) length

To assess and measure the effect of poly(A) tail length on POI mRNA translation, Four different mRNAs were generated encoding eGFP-HiBit and Myc-HiBit with poly(A) lengths of different sizes: 3, 40, 80 or 120 adenosines. All mRNAs were translated in parallel for a total of 120 minutes in the RRL system. The resulting luminescence readings were normalised to the average of the 120(A) sample per replicate (Figure 5A). Proteins were also detected by Western blotting (Figure 5B). All Poly(A) lengths tested achieved mRNA translation in the RRL system, however, an improvement of 40(A) over 120(A) for eGFP (p=0.006) and 40(A) and 3(A) over 80(A) for Myc (p= 0.0167, p=0.0365 respectively) was observed.

**Figure 5:**
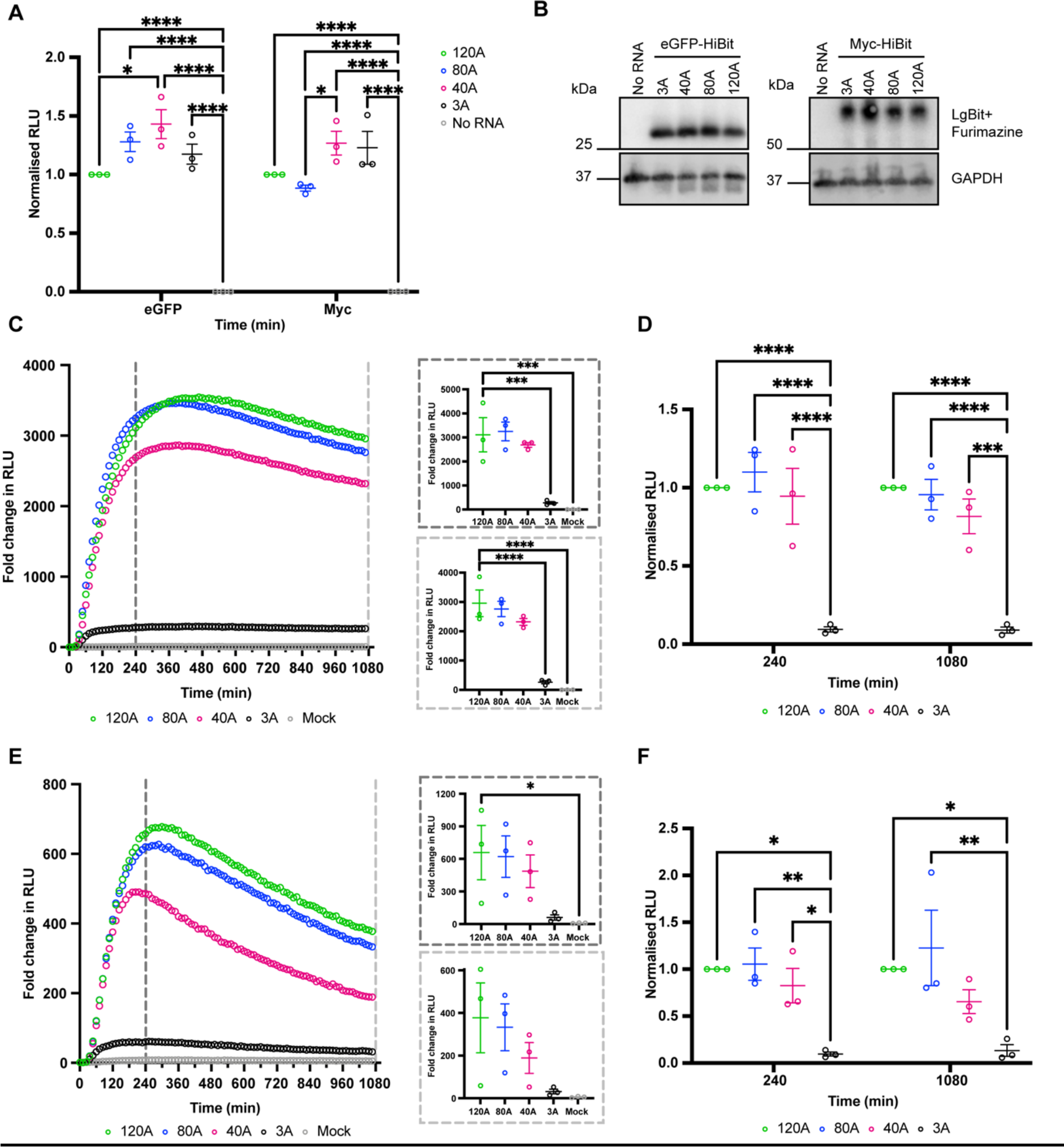
Effect of poly(A) length on HiBit-encoding mRNA translation is independent of the encoded POI. **(A)** mRNA encoding eGFP-HiBit or Myc-HiBit with poly(A) tails of 3 (black), 40 (magenta), 80 (blue), and 120 (green) adenosines was compared in the RRL system by incubating the HiBit mRNA for 120 minutes. Scatter plot depicting RLU of three biological replicates normalised to 120A eGFP-HiBit or Myc-HiBit, mean and SEM. **(B)** Western blot of pooled RRL samples with GAPDH as a loading control and eGFP-HiBit protein detected by LgBit and Furimazine. **(C)** Live cellular kinetic assay in HEK293 LgBit cells transfected with eGFP-HiBit with differing poly(A) lengths or Mock. RLU per sample was normalised to t0 to produce fold change in RLU; dots representing the mean of three biological replicates per time point. Mean and SEM at 4 hours (dark grey dashed line) and 18 hours (light grey dashed line) are shown, ordinary one-way ANOVA compared to 120A eGFP-HiBit. (**D)** Data at 4 hours and 8 hours was normalised to eGFP-HiBit 120A to depict changes in the HiBit signal and compared using Two-way ANOVA with matched values. **(E)** Live cellular kinetic assay in HEK293 LgBit cells transfected with Myc-HiBit with differing poly(A) lengths or Mock. Fold change in RLU from time 0 is depicted, with dots representing the mean of three biological replicates per time point. Mean and SEM at 4 hours (dark grey dashed line) and 18 hours (light grey dashed line) are shown, ordinary one-way ANOVA compared to 120A Myc-HiBit. (**F)** Data at 4 hours and 8 hours was normalised to 120A Myc-HiBit and compared using Two-way ANOVA with matched values. * p<0.05; ** p<0.005; **** p<0.0001.

Live kinetic recordings in HEK293 LgBit cells confirmed that the rapid translation of eGFP- HiBit post-transfection is common to all poly(A) forms except where the poly(A) tail is reduced to 3 adenosines (Figure 5C-D). In cells, the short 3A poly(A) almost ablated the translation of eGFP-HiBit (Figure 5C), the peak signal reached being significantly lower than that of any mRNA and 91% lower than 120A at 4 and 18 hours (p<0.0001, Figure 5D) and not significantly higher than that of the mock-transfected control at 4 and 18 hours (Figure 5C call-outs). The expression kinetics of mRNA encoding for Myc-HiBit recapitulated the observed impact of varying poly(A) lengths (Figure 5E-F). A poly(A)of 3 adenosines ablated Myc-HiBit expression to 90% at 4 hours (p=0.0104) and 87% at 18 hours compared to 120 adenosine tail (p=0.0141, Figure 5E).

## Discussion

The development of mRNA-based therapeutics has increased exponentially in recent years, and it has become clear that a crucial step in the development pipeline is mRNA sequence design and optimisation. A large effort has been devoted to the generation of computational tools to predict the optimal structure and composition of the RNA, with a particular focus on codon optimisation(8, 9), and the experimental validation has largely relied on existing assays which often employ reporter proteins (GFP and Luciferase) or the SARS-CoV-2 spike protein(7–9). In this work, we implement the HiBit tag in a pipeline for studying the kinetics of translation of exogenous mRNA *in vitro*, in the RRL system, and *in cellulo*, in HEK293 cells constitutively expressing LgBit. Importantly, the addition of the small component of the luciferase as a tag did not affect *in vitro* translational kinetics and offered a robust luminescent read-out specific to the tagged POI (Figure 1). Additionally, a luminescence but not a fluorescence read-out is compatible with the RRLs with a signal:noise that results in an assay with a Z’=0.8. The robust assay is compatible with applications in multi-well formats and, therefore faster and higher throughput than canonical assays for protein detection, which rely on antibody-mediated or radioactive labelling detection methods. The addition of the tag also permits the detection of the mRNA-driven expression of the POI over time in live cells constitutively expressing LgBit, treated with the luciferase substrate (Figures 2-5). The live luminescence recordings are vital in determining the expression window of the therapeutic mRNA, without the need to extrapolate information from reporter genes like engineered fluorophores or luciferases that translate very efficiently and are often stable proteins. In agreement, our data shows that the kinetics of expression between two POIs, eGFP-HiBit and Myc-HiBit are extremely different (Figure 3A-B). If no LgBit-expressing cells are available, assessment of POI-HiBit in cells can be carried out using the Nano-Glo HiBit lytic detection system (Promega), where cells are lysed and LgBit and the substrate are added to the lysates. The lytic method for detection of mRNA-encoded POI-HiBit has been recently employed for validation of an open-source computational mRNA optimisation tool(8), specifically for dose-response studies of mRNA administration at a set endpoint. The strategy is high-throughput and adaptable to different cell types but provides no information on the kinetics of expression of the administered mRNA. While time-course experiments can be carried out with the lytic system, it requires sampling of replicates at different time points instead of live monitoring of one same replicate over the time course of the assay, increasing the likelihood of variability between experimental replicates. Given the availability of techniques to generate stable cell lines, the engineering of LgBit knock-in cells in the model of interest unlocks the ability to conduct mRNA kinetic studies specific to the cell of interest and compare such kinetics between differing cell types.

To demonstrate the sensitivity of the mRNA-HiBit assay pipeline, we modified coding and non-coding elements of the mRNA and verified its ability to detect changes in mRNA translation. We confirmed the ability of both RRL and live cell assays to measure translation by comparing capped and uncapped mRNA, which only resulted in luminescence being recorded in the presence of the cap (Figure 2A-C, G-H). We next compared cap1 (CleanCap) to cap0 (ARCA) (Figure 2A-C). Significant translational benefit from cap1 has been seen in cells such as the murine immature dendritic cell line, JAWS II, and macrophages(60–62). We detected little difference in translation kinetics when comparing ARCA and Clean Cap in HEK293s (Figure 2G-H), suggesting changes in translation kinetics may be cell-type dependent(60, 61) and target cells should be evaluated. Outside of the scope of the RNA-HiBit assay, cap1 has been shown to reduce immunogenic response(63, 64) because 2’-O-methylation of the first nucleotide can help distinguish self RNA from non-self RNA by preventing recognition by RIGI and IFIT1 (Interferon-induced protein with tetratricopeptide repeats 1). Therefore, cap1 possesses an additional benefit of reducing immune silencing over the cap0 structure which is particularly attractive for *in vivo* use.

Engineering a hairpin structure within the 5’ of the mRNA significantly impacted the POI translation in cells, to the same extent of removal of the 5’UTR. Previous work suggests that mRNAs with limited thermodynamically stable secondary structures <-30 kcal/mol MFE) allow for efficient translation, whereas more stable secondary structures (>-50 kcal/mol) stall the 40S ribosome and decrease translation efficiency(16). In agreement, an mRNA with a 5’UTR containing limited thermodynamically stable secondary structures (∼–10 kcal/mol) was efficiently translated whereas translation was significantly attenuated with an mRNA containing a 5’UTR-Hairpin with an MFE close to -50 kcal/mol. These data support the fact that this assay can detect changes in translation resulting from different ranges of stability. Interestingly, the reduced translation was not observed in RRLs. Considering the nuclease-treated RRL has been depleted of all endogenous mRNAs, the assay does not recapitulate competition between different mRNAs for the ribosome and cap-poly(A) synergy(65, 66), which may account for the differences observed. These data highlight the need for in cell assays that allow discrimination of possible blocks to translation.

A key advantage of our live kinetic assay is its ability to measure both translation rates, in the early phases of the experiment, and protein turnover, after the initial burst of translation. To demonstrate this, we harnessed the highly regulated protein Myc and compared it to the expression of eGFP-HiBit (Figure 3A-B). Confirming our previous observation(49), the expression of eGFP in cells is far greater, and the signal is prolonged, unlike Myc expression which peaks at 4 hours and then declines rapidly. An advantage of using the HiBit system is that it enables the detection of the exogenously expressed POI, which can be difficult to detect if the endogenous equivalent is expressed in the cell line of interest. Indeed, no HiBit signal is recorded in mock transfection in favour of Myc-HiBit transfected HEK293 LgBit cells that do express abundant levels of endogenous Myc.

In a normal setting, the half-life of Myc is estimated to be around 20 to 40 mins(67, 68). Mutation of T58 to a non-phosphorylatable alanine (T58A) leads to an over 3-fold increase in Myc half-life(67) which promotes growth and proliferation(69) and is a mutational hotspot in lymphomas(70, 71). A second phosphodegron, T244, has also been identified, but less is known regarding its ability to stabilise Myc protein(55). In our HiBit assays, all phosphoablation mutants translated equally in the RRL system. In HEK293 LgBit, only Myc-T58A-HiBit, displayed increased expression over the WT Myc-HiBit. Interestingly, this was true at the peak of expression when T58A resulted in a 42% increase in Myc-HiBit signal at 4 hours, which was not sustained over time. At 4 hours the translation rate is in the exponential phase and likely greater than the degradation rate, therefore the effect of changes in protein stability can be observed, as confirmed by proteasomal inhibition (Figure 3F-G). Importantly, if these studies had been conducted as a single end-point evaluation at 18 hours, all versions of Myc would have been deemed comparable, despite the known phenotypic differences that Myc-T58A triggers(69–71). Since other regulatory elements that dictate Myc turnover exist(70, 72, 73), further sequence modifications were employed to determine whether a more stable translational output was achievable. Applying a published, open-source algorithm(9) to the best-performing stability mutant (T58A), codon optimisation and mRNA stability strategies were implemented and tested using the HiBit assay system. In HEK293 cells, translation of Myc T58A-HiBit was greater for a CAI-optimised sequence, producing a delayed peak and prolonged translation with an ∼80% increase in signal 18h post-transfection compared to non-optimised WT and T58A Myc-HiBit. Contrarily, MFE optimisation led to reduced translation with a resulting 70% reduction of Myc-HiBit signal. However, MFE optimisation did lead to a more stable, albeit lower, Myc-HiBit signal, within the timeframe of the described experiment. These observations agree with previous reports that highly structured RNA sequences can enhance mRNA stability in cells, but the highly structured RNA decreases the ability of the cellular translation machinery to process the RNA(7). Interestingly, both optimisation strategies combined were insufficient to bring Myc close to the expression levels and steady state of protein synthesis and degradation that can be sustained over time, as seen with eGFP. Together these findings highlight how the optimisation strategy should be tailored to the encoded POI and its biological output. Overall, our data supports the ability of this HiBit assay system to report on changes to the coding sequence, both with regards to improving mRNA translatability through codon optimisation strategies and protein engineering through stability mutants.

An essential aspect of the development of mRNA-based therapeutics is the incorporation of modified nucleotides, which are chosen based on their effect on the translation efficiency of the mRNA and their ability to reduce immunogenic responses(74–76). Importantly, our data demonstrate that the choice of modification will depend on the encoded gene and therefore modified nucleotides should be tested on the therapeutic mRNA, not a reporter protein. By comparing modified nucleotides incorporated in mRNA encoding eGFP-HiBit and Myc-HiBit, we found the choice of modifications only minimally affected eGFP-HiBit expression but greatly affected Myc-HiBit translation, with incorporation of pU resulting in a higher and more sustained expression over time. It is important to note that the data presented in this work does not address how the different modifications may affect translation rates in alternate cell types. By generating stable cell lines expressing LgBit in target cell, the HiBit system would be valuable to enable comparison to be made.

Finally, we assessed the contribution of differing poly(A) lengths to mRNA translation *in vitro* and *in cellulo*. In the RRL system, no advantage of increased Poly(A) length was measured, when ARCA cap is present (Figure 5 A-B), but loss of cap could not be rescued by the addition of a long (120 adenosine) poly(A) tail (Figure 2 A-C). Therefore, for protein synthesis applications, a tail with as little as three adenosines is sufficient for efficient translation of capped mRNA. This is in agreement with previous reports which demonstrated that in nuclease-treated RRLs, such as the Flexi Rabbit reticulocyte lysate kit, poly(A) tail length had minimal effects on the translation of capped mRNA(77, 78). In cells, minimal translation was recorded from constructs containing mRNA with a poly(A) 3 adenosine long. Per the literature, high-affinity binding of PABPC to poly(A) tails requires approximately 12 nucleotides, with a bound PABPC covering around 30 adenosines in length(32, 79), meaning an mRNA with 40A tail will only be bound by one PABC protein versus two or more which have been shown to increase translation efficiency(80–82). mRNA encoding eGFP-HiBit or Myc-HiBit with a poly(A) tail of at least 40A was efficiently translated likely due to the binding of a PABPC molecule. No difference was detected between the translation of mRNAs with a tail permissive of binding of two (80A) or more (120A) PABCP, indicating that for mRNA therapeutic design, 80A tails are sufficient. These data demonstrate the ability of the HiBit assay pipeline to detect translation rate differences brought about by differing poly(A) lengths which recapitulate published data.

Overall, our findings demonstrate that the high-throughput HiBit pipeline that we describe facilitates mRNA translation measurement both *in vitro* and *in cellulo*. While the *in vitro* translation system of the RRL yields detectable translation changes by coding and non-coding elements of an exogenous mRNA, it is not a reliable predictor of efficient mRNA translation in live cells, possibly due to discrepancies between model systems, but can prove useful as a first-line screening tool. Implementing a live cell time course assay, as we describe, overcomes this limitation and unlocks essential information on the dynamics of protein synthesis and degradation of a POI encoded by an exogenous mRNA. Importantly, the live kinetic assay can record differing windows of peak expression of a therapeutic that will depend on the encoded POI and therefore can inform on when phenotypic output could be expected and temporal windows in which modifications could improve the expression of the therapeutic.

## Data Availability

Further information and requests for resources and reagents should be directed to, and will be fulfilled, by Dr Catherine Wilson and Dr Camilla Ascanelli.

## Acknowledgements

The authors thank Laura Itzhaki for the LgBit HEK293 cell line and Annabel Cardno for sharing her expertise on HiBit.

